# Evaluating host to host transplants as a method to study plant bacterial assembly

**DOI:** 10.1101/2021.08.30.458296

**Authors:** Christopher Baldock, Neil Wilson, Rosalind Deaker

## Abstract

The ability to predict plant microbiome assembly will enable new bacterial-based technologies for agriculture. A major step towards this is quantifying the roles of ecological processes on community assembly. This is challenging, in part because individual plants are colonised by different communities of soil bacteria and it is difficult to estimate if the absence of a given species was a) because it was not present in the soil to colonise a given plant or b) it went locally extinct from competition, predation or similar. To minimise this uncertainty, the authors develop a mesocosm system to study bacterial communities of individual plants by replicated transplantation to a recipient host plant population, ensuring new hosts receive a homogenous species pool for colonisation. We sought to understand which factors affected the transplant and, what the main drivers of variation in the model communities were. A nested factorial design was used to investigate the transplantation of cultured or total, root or leaf associated bacterial communities from donor host species to surrogate host species. Specific metrics were developed to quantify colonisation efficiency of communities. The results show the root communities were more effectively transplanted than leaf communities, with leaf communities more susceptible to contamination. For root communities the strongest driver of beta diversity was the donor host species, and for leaves it was the surrogate host species. Overall, the results reveal that root, but not leaf communities are amenable to transplant reflecting their differing ecological drivers. This work provides the basis to develop a plant microbiome transplant system.

## Introduction

Predicting patterns of microbial community assembly is critical to their manipulation in crop production(1). Understanding the rules of community assembly and what makes a community resilient will be essential to the effective delivery and persistence of bacterial inoculants, consortia or synthetic communities (2). Selection pressures from host factors, abiotic variables and interspecies interactions can all theoretically drive the structure and variation in communities (3). There is significant appreciation of how host factors and environmental variables shape root and leaf bacterial communities. From specific taxa that are heritable in the maize rhizosphere, to the enrichment of actinobacteria and firmicutes in rhizosphere and root microbiomes under drought conditions (4–8). However, how these drivers interact with bacterial-bacterial interactions remain understudied. Advanced understanding of bacterial dynamics in the human gut microbiome has been facilitated by periodic sampling from the same host, allowing inferences to be made from longitudinal data on the same set of species in the same host (9). Destructive sampling means this is not possible in plants and one must use a population of hosts to determine statistical probabilities of community dynamics.

Root and leaf communities are colonised by soil microbes (10–12). Thus, soils with high turnover in community composition will result in less overlap in the species pool that can colonise each host plant. It can be difficult tease apart the turnover in soil communities from other drivers of plant microbiome beta diversity, such as host genotype or species competition. The effect of different species pools has been demonstrated to inflate beta diversity in macro ecological systems and may confound the effect of other selection pressures (13). It also creates uncertainty in estimating interactions from spatial samples. This is because it is difficult to know if the absence of a species from a particular host is the result a competitive/predatory interactions or because it was not present in the species pool to begin. One may consider a census of the soil proximal to each host plant; however, gaining sufficient sequencing coverage to have high certainty in taxa detection is costly, and, modelling this is difficult (14).

An alternate way to control for this is to use a pre-defined species pool that colonises the plant. This can be done through a gnotobiotic system with a synthetic community (SynCom) of bacteria (15). However, obtaining representative bacterial collections to perform such experiments may be beyond the scope and/or resources of many research groups. Moreover, this system lacks flexibility as it is generally restricted to core or readily culturable taxa. This limitation may miss emergent properties of particular sets of organisms seen *in situ*.

Here, a mesocosm system was developed and tested to study quasi-natural bacterial communities. Inspired by analogous studies of the gut microbiome, whereby human gut bacterial communities were sampled *en masse* and transplanted to surrogate, gnotobiotic mice(16). In this study microbial communities were transplanted from donor plants into surrogate plants in mesocosm system that is autoclaved but not gnotobiotic. This approach allows the separation of soil communities edaphic properties, environmental conditions and the plant and associated communities, enabling researchers to ask critical ecological questions.

The aims of this study were to determine whether a plant-host microbiome can be successfully transplanted in the mesocosm system, which factors affect the transplant and what drives the variation in the model communities.

## Methods

### Experimental design

The experimental approached used a factorial design to investigate the effect of root-to-root or leaf-to-leaf community transplants (Figure S1). The root and leaf samples were sourced from commercial avocado and banana farms in Northern Queensland, Australia. These species were chosen for study because they have high commercial significance in horticulture, but, their interactions with microbial communities can’t be studied easily in laboratory settings as they are large perennial species that are not feasible to grow in these conditions. The communities were transplanted as a homogenate, (total community) or from an *en masse* cultured community, to ryegrass or tomato surrogate plants in control laboratory settings. These plants were chosen to represent a monocot and dicot species and because they are annual species amendable to growth in laboratory conditions. Kaolin clay was used as a synthetic soil and Murashige and Skoog media (MS) was used for the supply nutrients to keep environmental conditions constant.

Sampling occurred with gloves and 70% ethanol cleaned utensils, before shipping on ice to the University of Sydney. On arrival, samples were homogenised with PBS (10ml for every 1 gm of plant tissue), in an ethanol-sanitised blender, before being frozen at minus -80°C in a (30% v/v) glycerol stock to plant homogenate until use.

An alternative to magenta boxes (such those used by (15)) was sought because they are costly for large sample sizes and are difficult to scale to larger plant species. A system was devised of pots (Danbar, dimensions 50×70×50 mm (W X H X L)) enclosed in sterile plastic (200 mL Twirl-Pak bags) to prevent cross contamination. Pots were filled with 110 mL of autoclaved synthetic soil (30:70 volume ratio of Boral builders clay to washed river sand). To this, 20 mL of ½-strength MS and 10 mL PBS was added.

Seeds (Mackay annual ryegrass or yates tiny tom) were surfaced sterilised in 4% bleach for 5 mins and then rinsed seven times with MQ water. Either five tomato seeds or approximately ten ryegrass seeds were sown in each pot. The difference in seed quantity between species was to compensate for the lower leaf biomass of ryegrass. Forty containers were placed in a tray, representing all the samples to be inoculated by the same donor species (i.e. cultured and uncultured).

### Inoculation and growth conditions

For experiment one surrogate plants were inoculated with the plant homogenate or a cultivated version of the donor community. To obtain the plant homogenate the donor plants were homogenised with phosphate-buffered solution in a Warrick blender and 1 mL of 1×10^−1.5^ diluted extracts was inoculated to each container. This was chosen to allow enough inoculum to transplant to a sufficient number of surrogate plants, but as not to reduce the population density too greatly. The estimated CFU/mL of the uncultivated fraction were ∼ 1×10^6^. For the cultured fractions the plant homogenate was plated at high density (1×10^−1^) on eight R2A agar plates. The plates were harvested by scraping into 9 mL phosphate buffered solution (PBS) and then diluted 10-fold by transferring 1 mL to 9 mL PBS (i.e. 10^−2^); One mL of this was pipetted into each pot.

Three days after planting, donor communities were transplanted to seedlings/seeds. The leaf and root communities were transplanted in the same manner by inoculating the soil with 1 mL of the transplant community. After inoculation a second A4 plastic sleeve was placed around each sample to minimise contact and dispersal between communities. This was in addition to the Whirl-pak bag mentioned above. Pots were randomised but each tray had a different donor community and compartment.

Next, trays were placed two abreast under two lighting platforms. Tray orientations were shuffled every two days to ensure plants received even lighting. Additionally, alternate trays from each platform were swapped at the same time to control for the lighting difference between the platforms. Plants were watered with MQ water every second day (excluding weekends).

### Sample Harvesting and processing

For harvesting, ethanol-cleaned scissors were used to remove the stem from the roots. Leaf samples were transferred to a sterile 50ml falcon tube with three 20mm glass beads sterilised prior using 1.0 % sodium hypochlorite for 10 min followed by washing in sterile MQ water. For roots, forceps were used to shake away excess soil and rhizosphere soil was removed with three washes of MQ water in a 50ml falcon tube (vortexed 2 × 20s each wash). Roots were transferred to a 50 ml Falcon tube containing sterile glass beads. All samples were frozen at -20 for no longer than four weeks.

### DNA extraction and Illumina library preparation

Sample tubes were frozen with liquid nitrogen and homogenised on a vortex to a powder. To this, 3 ml of lysis buffer 1 (0.58 g NaH_2_PO_4_.H_2_O, 1.55 g Na_2_HPO_4_.7H_2_0, MQ H_2_O to 100 ml) was added before vortexing and aliquoting 700μL volumes into 2ml bead tubes with a 100μL of silica carbide beads. A 125 μL volume of Lysis buffer 2 (4 g SDS, 6mL 0.5 M EDTA pH 8.0, 15 mL 1 M Tris-HCl pH 8.0 and MQ H_2_O to 50 ml) was added and bead beated for 45s at 2000rpm on a Mobio homogeniser. The homogenate was centrifuged for 2 min at 10000 prm and then and 700μL of supernatant was transferred to a new tube with 125μL K acetate (7.5 M). This was incubated for 5 min before being centrifuged at 10000rpm for 5 mins. From here, samples were purified using Ampure XP magnetic beads before eluting into 1/10 TE buffer.

The V3-V4 region of the 16s rRNA gene (17) was sequenced on the Paired 2×300bp illumina Miseq V3 chemistry. A dual index Illumina library preparation with PNA clamps was used (18,19). In a 96 well format, PCRs consisted of an average 6ng template DNA (+- 2 ng), 2ΜL of MyTaq red 5X reaction buffer, Mytaq DNA polymerase (0.1 μL), and 0.5μL of each 16S primer. The first thermal cycling protocol ran as follows: 95 degrees C for 3 min, followed by 26 cycles of 95 degrees for 20s, 78 degrees for 10s, 55 degrees for 20s, and 72 degrees for 30s, and finally by 72 degrees for 10 min then cooling to 4 degrees. This reaction was performed in triplicate, before being pooled and diluted 1:100 in Millipore water. A second indexing PCR was carried out in a 20ul volume (mix details), at: 95 °C for 3 min, followed by 10 cycles of 95 °C for 20s, 60 °C for 20s, and 72 °C for 45s, and finally by 72 °C for 10 min then cooling to 4 °C,

Samples were quantified with Quantiflour DS DNA kit (New England bio labs) on a Biotek Synergy H1 plate reader, then pooled at equimolar concentrations, before being purified with Ampure XP beads and diluted to the correct molar concentration

### Bioinformatics, Statistical Analysis and Data Visualisation

Sequences were demultiplexed with DeML(20). All subsequent processing and analysis were carried out in R version 3.5.1 (21). Sequence variants and taxonomic inference was done in the DADA2 package (22). Exploratory analysis was done in phyloseq (23). Generalised linear latent variable models were fit with BORAL (24) and visualisation done with ggplot2 (25). ANOVA was carried out in base R and using the “car” package to calculate type three sum of squares (26). The R code to reproduce this analysis is available at (https://github.com/ch16S/host_to_host_transplant).

## Results

### Transplant efficiency differs between donor and recipient host, compartment and culturing

This study transplanted communities from donor (banana or avocado) to recipient (tomato or ryegrass) plants. Transplantation was done root to root or leaf to leaf. It was sought to understand if these communities could successfully be transplanted and if the donor or recipient plant impacted the composition of the transplant. Moreover, propagating a community may be useful to study over multiple experiments. Thus prior to transplant, communities were cultured *en masse*, or, transplanted as a leaf or root homogenate (referred to as “total”). How the transplanted communities change over time was also examined. The design of this experiment is useful to understand how taxonomic composition is affected, but not how the transplanted community’s functionality is affected, although the latter could also be achieved.

The 16S V3-V4 variable region of each transplanted and donor community were sequenced. This enabled the origin of taxa in the transplanted communities to be identified. The taxa could be from one of three sources: the target donor plant (target donor), from another donor plant (nontarget donor) or from the environment (random). We deem target donor taxa to be the fraction of the community that was used to inoculate the recipient plant e.g. cultured banana root community (Figure 1, top image). Non-target donor taxa are from a donor community, but not the one the recipient plant was inoculated with (Figure 1, middle image). The third group were taxa not detected in any of the donor communities, these were considered random taxa (Figure 1, bottom image). They were likely indigenous to the plant-growth environment.

**Figure 1.**
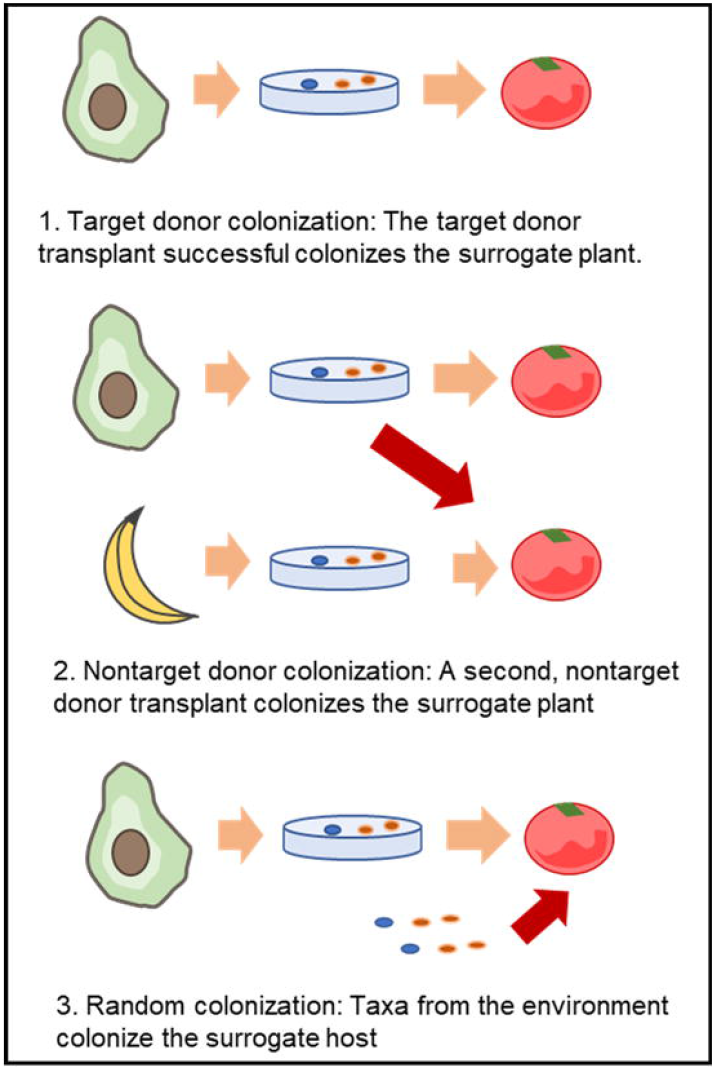
The three possible sources of taxa that colonise the surrogate plant hosts. The avocado and banana represent the donor plants while the tomato represents the recipient host. The petri dish represents the transplant communities and the orange arrow represents the desired colonisation source of the surrogate host, while the red arrow represents a contaminate or not target source of colonisation.

Here, a transplanted community with high amounts of target-donor taxa and, low amounts of non-target and undetected taxa are considered to have a high transplant success. We first visualised the proportion of reads from each of these groups (Figure 2a). The leaf communities had the lowest target reads. Ryegrass leaf communities had a higher proportion of nontarget reads while tomato had higher fractions of undetected/random reads. For the root samples cultured communities had a higher proportion of nontarget reads and total communities had a higher proportion of target reads. All communities were variable in their recovery of the total starting community (Figure 2b). The main trend noted was that communities transplanted to ryegrass tended to have higher recovery than ones transplanted to tomato plants. Interestingly, the communities with highest proportion of target taxa had the lowest species recovery of their donor community.

**Figure 2.**
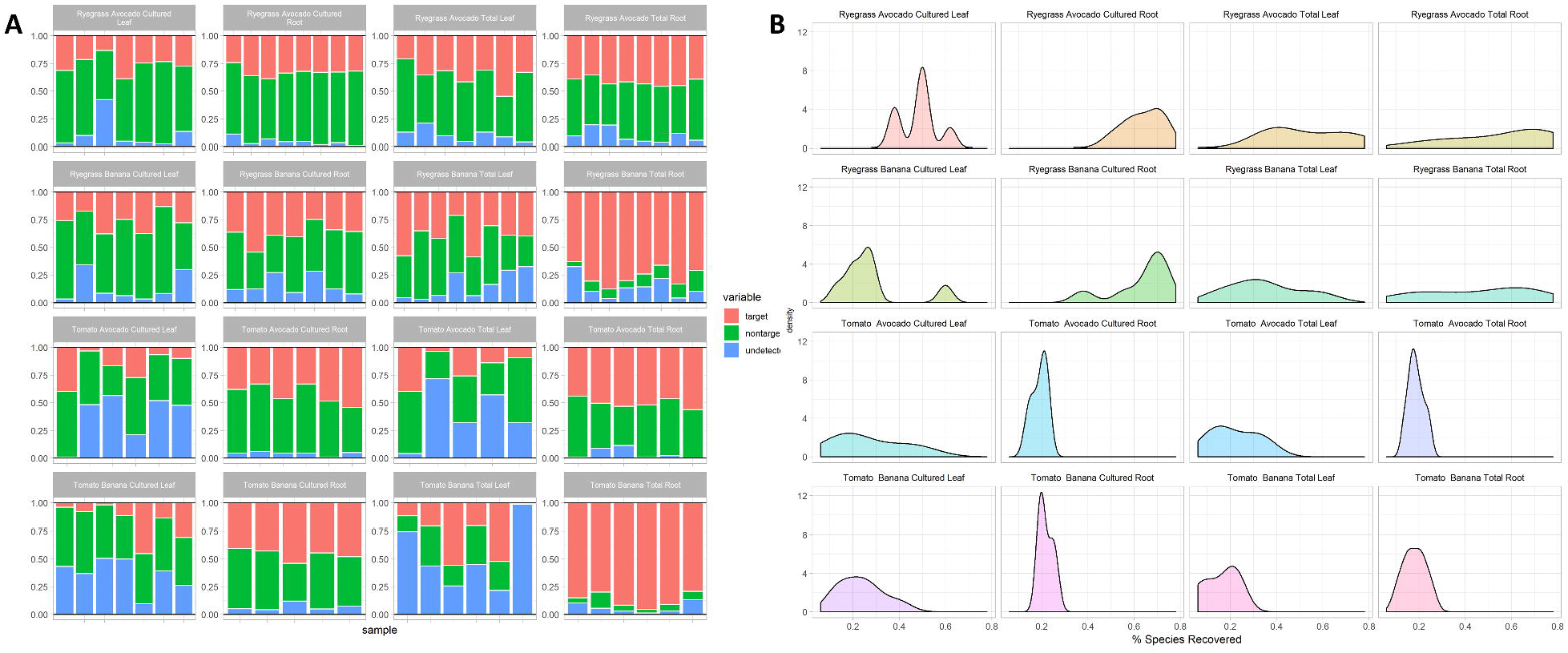
A) The relative abundance of the three species pools: target taxa(red), non-target taxa (green) and random taxa (blue) the titles give the combination donor and recipient plant compartment and whether the transplant was cultured. B) The smooth kernel density distribution of the taxa recovered (% out of 1) in each donor community.

The absolute differences in colonisation between these three species pools couldn’t be assessed as the sequencing data is in the simplex (27). Instead, the change in pairwise log-ratios of between the three groups (target, non-target and random) were quantified to represent their relative colonisation (28).

Five measures were developed to assess transplant success. The measures were was based on binning the reads into the aforementioned three groups (target donor, non-target donor and random), after agglomerating to genus level taxonomy.

The first metric quantified the log ratio between the total abundance of the target-donor taxa to the total abundance of non-target donor taxa (called the nottarget abundance ratio or NTAR, Figure 3, row 2). The second measure quantified the log ratio of the target donor abundance to the total abundance of random taxa (called the undetected abundance ratio or UDAR, Figure 3 row 2). For these two measures, if the value was greater than zero there was a higher proportion of target taxa to that of nontarget or random taxa. If the value was below zero, then there was a lower proportion target taxa. A successful transplant will have a positive value for both measures, as this shows the target taxa make up the majority of the community.

**Figure 3.**
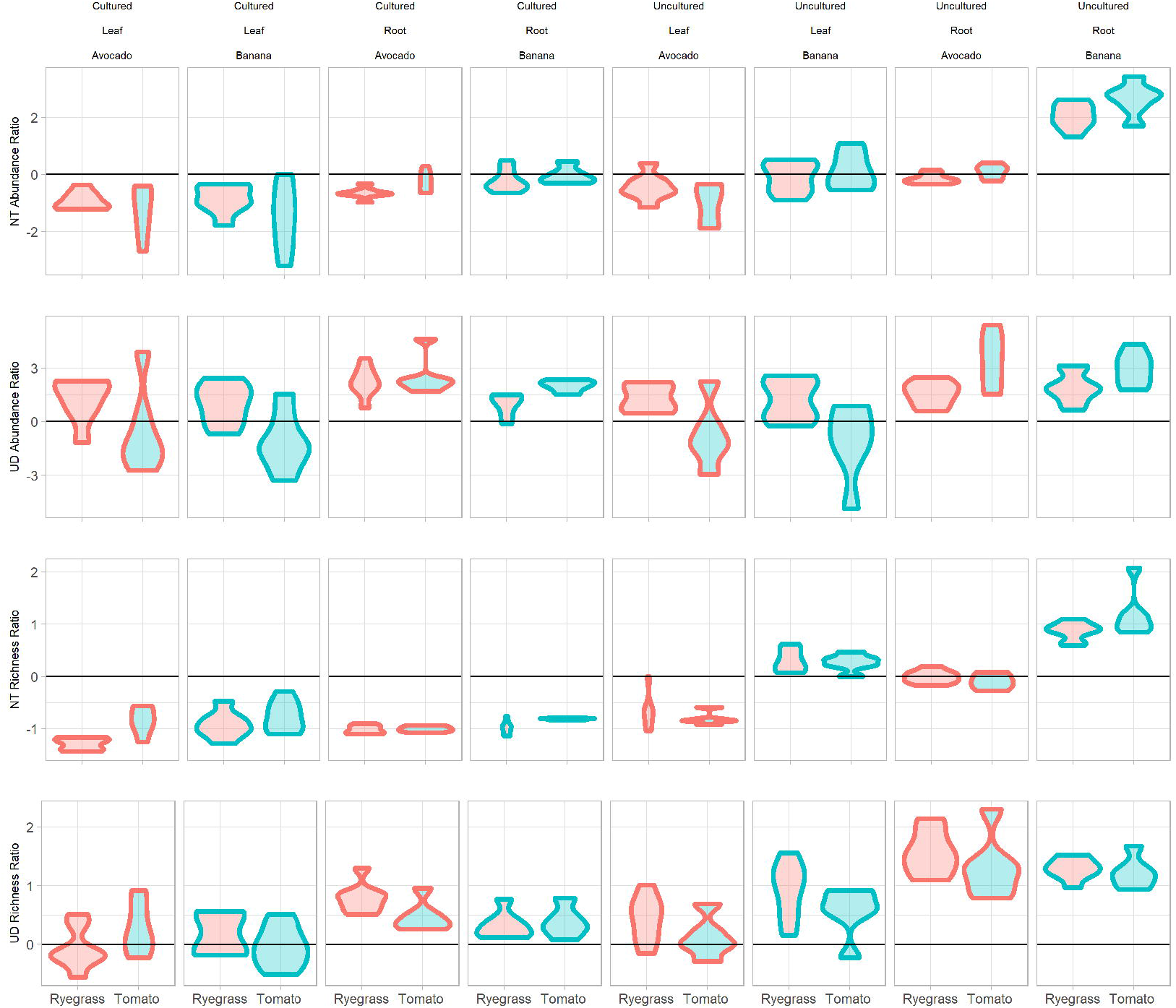
Violin plots of the metrics across the eight donor communities. Each row represents a different measure. Values greater than zero indicate there is more reads or species belong to the target group than then non target or random groups while values below zero indicates the opposite. Top row: The log abundance ratio of non-target taxa. Second row is the random/undetected abundance ratio. Row three: non-target richness ratio and row four: The undetected/random richness ratio. Red fill indicates ryegrass recipient plants and blue fill indicates tomato recipient plants. The black line indicates zero, where there is equal proportion of the two groups being compared. Note the violin plot shows a mirrored empirical probability distribution of that data.

These measures do not account for the species richness. This can be misleading if a few dominant taxa inflate the proportion of reads from the donor community or vice versa. Thus, the log-ratio approach was extended to quantify ratio of species richness of the target-donor taxa to the non-target (called the not-target richness ratio or NTRR, Figure 3 row 3) or random taxa (called the undetected richness ratio or UDRR, Figure 3, row 4). The species richness ratios are interpreted in the same manner as the abundance ratios.

Lastly, to assess how much of the donor community was successfully transplanted, the percentage of the target-donor taxa recovered in the surrogate community was calculated (Figure 2b).

For the aforementioned plots in figure 3 each column is a different donor community. The violin plots in the left of the pane show the tomato communities and the right side show the ryegrass communities. The first row shows the non-target abundance ratio, the second row shows diversity ratios undetected or random abundance ratio, and third and fourth showed the nontarget and random richness ratios respectively.

To understand which factors had the biggest influence of transplant efficiency the five metrics were modelled as an ANOVA with the factorial design as explanatory variables. The model included donor species, surrogate plant host (plant), plant age (time), root or leaf (compartment), whether the total or cultured community was transplanted (cultivated uncultivated) as well as their two- and three -way interactions. The effect sizes (ETA^2^) are summarised in Figure 4 and Supplementary table 1.

**Figure 4.**
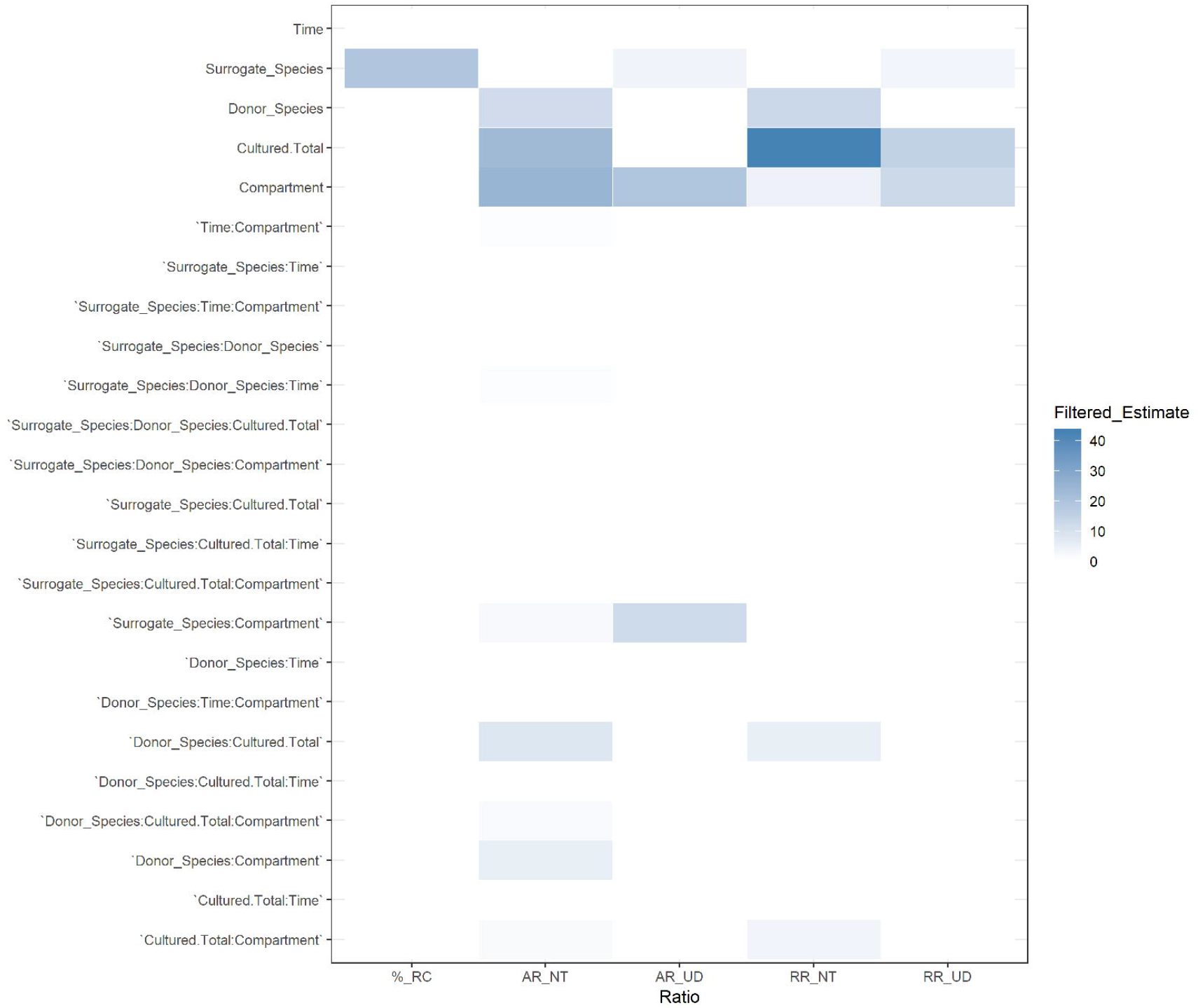
Heatmap of ANOVA (Type 3 sum of squares) effect sizes for each metric using ETA^2^. Values are only for significant terms (p<0.05). The X-axis indicates the five measures. %_RC = taxa from the original community recovered in the surrogate one, AR_NT abundance = non-target abundance ratio, AR_UD undetected abundance ratio, RR_NT non-target richness ratio and RR_UD is the undetected richness ratio. The factors and their interactions are shown in the y axis. Compartment is root or leaf, cultured.Total is if the transplant was cultivated *en masse* first or not, donor species is donor plant species, surrogate plant species is the recipient host and time is time point (4 or 6 six weeks). A “:” indicates an interaction term.

The ANOVA showed compartment to be the most common main effects and involved in the most interactions. This can be visualised in figure 3 and table S1.

The number of taxa recovered from the non-target donor species was significantly affected by surrogate host with tomato plants showing lower recovery than ryegrass, as seen in figure 2B. The not-target abundance and nontarget richness ratios displayed similar trends in significant model effects (Figure 3, row 1 and 3, Figure 4). This involved significant main effects for compartment, cultured/total and donor species as well interactions between these terms and the recipient host species. The factor Cultured/total in the nontarget richness ratio explained the most variation of any model, with *en masse* cultured communities more susceptible to colonisation by other donor communities (Figure3, row 3). Overall recipient hosts received total community transplants scored better than cultured ones, and root transplants performed better than leaf transplants.

For UD ratios, i.e. random taxa, there was fewer significant terms than the nontarget ratios. The undetected abundance ratio had significant terms for surrogate species, compartment and their interactions (Figure 4). For the UD richness ratio there was significant terms for cultured/total, surrogate species and compartment. Generally, leaf transplant had higher richness and abundance of random taxa, and tomato recipient plants had more random taxa than ryegrass (Figure 3, row 2 and row 4). Finally, for the percentage species recovered the only significant term was surrogate species, with ryegrass recovering more transplanted species than tomato.

Based on the metrics the uncultivated banana root and leaf donor communities were transplanted most efficiently (Figure 3, column 8). The avocado and banana cultured transplants performed the least efficiently in these five metrics. The cultured banana root and leaf communities had the highest recovery of donor taxa. In sum the banana communities were amendable to transplant than avocado and the root communities more amendable than the leaf. Across all samples there was significant amount of cross-contamination. The root had more non-target taxa than the leaf, and the leaf more random taxa than the roots.

### The beta diversity of transplant communities

The colonisation metrics showed the donor communities colonised the surrogate plants at different efficiencies and that contamination was an issue for all communities. It was next sought to understand how this variable colonisation efficiency shaped surrogate community assembly and whether patterns of beta-diversity could provide insight into the source of cross-contamination. To investigate this, the sequence variant table of the leaf and root surrogate communities were modelled with generalised linear latent variable models (GLLVMs). The latent variables (three in this instance) find unknown axes of variation in the data, that can be used analogously to principal components. They also enable joint modelling of the taxa distribution to account for correlation between them. A negative binomial with a log link function was used to model the counts while controlling for overdispersion from the mean-variance relationship. A fixed row-effect was included to control for sequencing depth. Two types of GLLVMs were fitted; a GLLVM with no covariates (pure latent variable or PLV), as well as a correlated response (CR) GLLVM. The PLV is akin to an unconstrained ordination, whereas the CR accounts for covariates. If the CR model explains the data well, patterns seen in the ordination from the PLV will disappear. Separate models were run on SV tables collected from the roots and leaf surrogate communities. The GLLVMs were fit as Bayesian models in the R package “BORAL” (24).

The latent variables of the GLLVMs can be seen in Figure 5. The PLV-GLLVM of the root samples shows clustering as follows: donor species on the first axis, surrogate host species on the second axis and cultivation and time on the third axis (Figure 5A). For surrogate root CR-GLLVM the experimental design fit the data reasonably well. The CR ordination does not show the clustering of the PLV (Figure 5B), indicating the experimental design explains much of variation in the PLV.

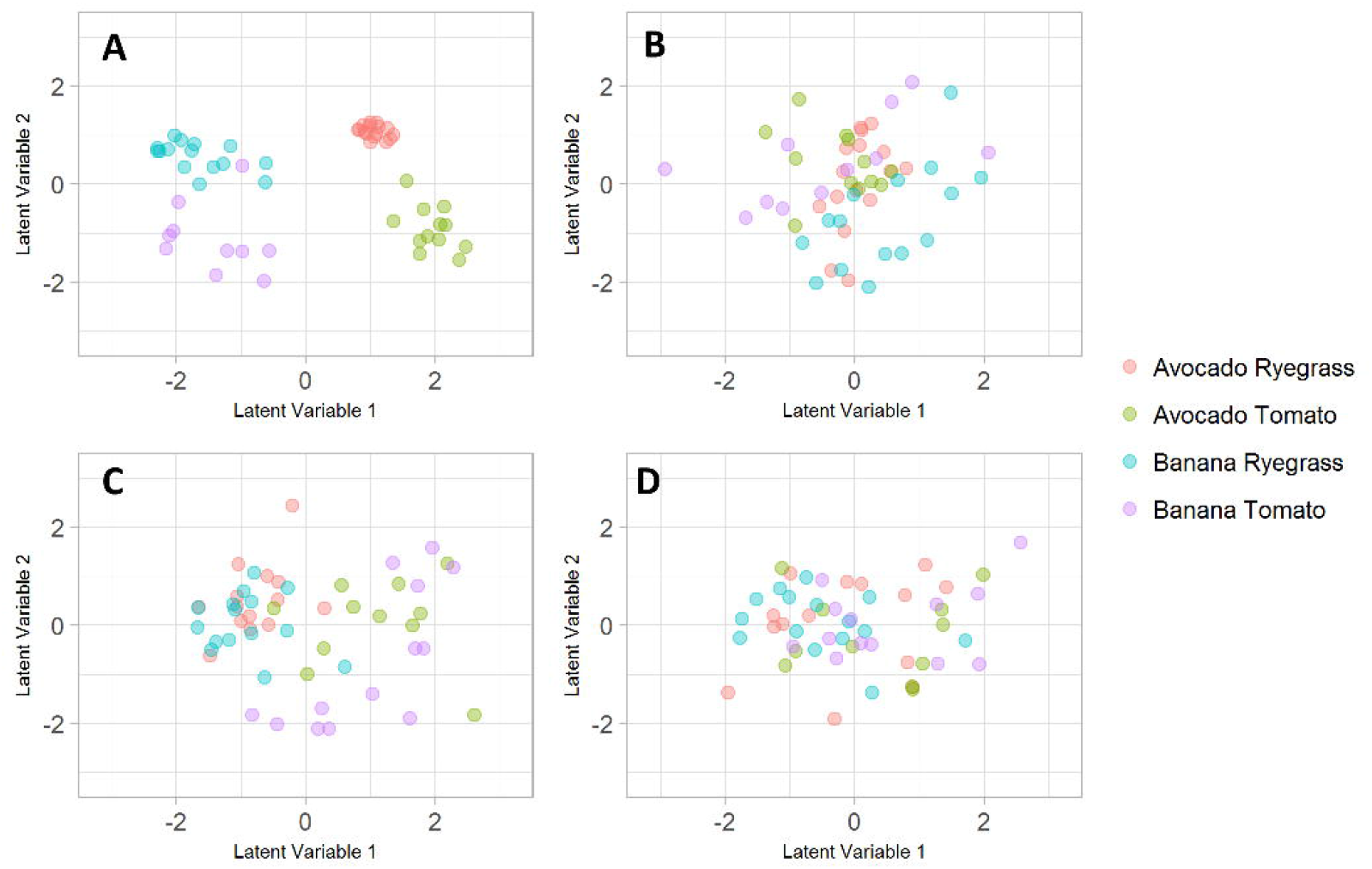

The leaf PLV-GLLVM showed less distinct clustering than the root PLV-GLLVM (Figure 5 C). The first axis displays a tight cluster of ryegrass communities, separated from tomato communities. The second axis separates the banana tomato communities from the rest. The CR-GLLVM, fit with full experimental design, removes some clustering but there is still a lot variation in the leaf CR-GLLVM ordination Figure (5 D). This corroborates the earlier findings that leaf colonisation was much more variable and indicates that factors than the experiment design are driving variation in the leaf surrogate communities

Figure 4 Latent variable ordinations. A: a pure latent variable model fit to the root data, B: a correlated response latent variable model of the root data (with all factors as explanatory variables). C: a pure latent variable model fit to the leaf data. D: A correlated response model of the leaf data (with all factors as explanatory variables). Legend indicates the surrogate plant species and the donor community used.

To understand what factors explained the species distribution in the CR-GLLVMs, the coefficients were plotted for each taxa from the root and leaf models. Figure S2 shows the proportion of taxa whose coefficients are significant for each model term, where significant means the 95% credible region does not overlap zero. The most influential factor is donor species (i.e. avocado and banana) for surrogate root communities (35% taxa affected) and host plant (i.e. tomato or ryegrass) for surrogate leaf communities (∼14% of taxa affected).

## Discussion

This set of experiments evaluated a plant mesocosm system for transplanting *in situ* communities. Earlier studies have successfully transplanted bulk, rhizosphere and foliar communities (29–32). To the author’s knowledge this is the first study to develop specific metrics to evaluate the transplant efficiency of donor communities.

Root and leaf habitats are known to exhibit different patterns of diversity (11,33). This study sought to understand if leaf and root differed in their amenability to transplant. Its shows that irrespective of donor or surrogate host species, the transplantation of a leaf community is more variable and less efficient than that of a root community. The leaf communities were driven by recipient host species more than donor species. This is in agreement with (33) that found leaf communities were driven by genotype more than root communities in *Boeshea stricta*. It may also reflects the disparity between the colonisation technique used here and the real-world colonisation pathways. Here the donor community was transplanted to soil, which would go on to colonise the leaf. However, in real settings, the leaf is colonised by soil, air and neighbouring plants (15,34,35). Bai et al. (15) used a spray technique to colonise leaves and found it provided different communities than when inoculating the soils in SynCom experiments.

Both root and leaf samples had contamination but from different sources. The surrogate leaf samples had higher proportions of random taxa (Figure 2), and more variation in their colonisation efficiency (Figure 3) and beta-diversity (Figure 5) than the root surrogate samples. As leaf communities were exposed to the growth room air it likely there was more chance of aerial colonisation by environmental taxa which may explain the lower UDRR and UDAR scores.

The roots samples had lower proportions of random taxa, but high proportions of non target taxa particularly in hosts that received cultured transplants. The cultured transplants clustered with the total community transplants of the same donor species on beta diversity ordinations (Figure 4). This pattern indicates the source of the contamination in surrogates root samples transplanted with cultured communities was their total community counterpart e.g. total avocado root contaminated cultured avocado root transplanted samples. If the contamination was from indigenous microbes, like the leaf communties, one greater dispersion and less clustering on the ordination plots.

The results here show that the transplant system in its current form requires optimisation. Aerosol/soil contaminants successfully colonise non-target host plants to reach high relative abundances, presumably from low initial densities compared to the target inoculant (Figure 2). This suggests that such bacteria had high relative fitness for a particular niche not filled by the taxa in the transplant community. Following this future work could extend the system to use more communities and to create more niches by modifying the soil chemical and physical properties and by applying stressors to the plant (e.g. drought) to study the interplay between plant host status, environmental and bacterial interactions. This would enable researchers to disentangle the drivers of community assemble and could be considered as an alternative method to artificial microbiome selection (29,36)

Interestingly, banana derived communities colonised the roots and leaves more effectively than communities from avocados across all metrics used, and this was irrespective of donor plant type. This suggests that some community level property may at play, perhaps related to metabolic complementarity or less antagonistic tendencies (37,38). Studies have used overlapping sets of bacteria to generate synCom in highly controlled gnotobiotic systems to study their interactions and phenotypic effect on plants (39,40). The results here raise the possibility that such an approach could be extended to transplanted communities. This could involve screening dozens of transplanted communities in the plant mesocosm system. The results could be used to identify taxonomic or functional groups associated with colonisation success.

In conclusion, the mesocosm developed here shows some potential, although it had variable success at transplanting plant associated communities. The leaf and root communities were influenced by different factors. The root communities were driven by their donor plant species, whereas the leaf was driven by the host plant and was generally more stochastic. This correlated to differences in transplant efficiency. Leaf communities had poor transplant efficiency and were susceptible to contamination from the environment. Root communities had greater transplant efficiency but also had higher cross contamination from other donor communities. Overall this system enables a new understanding of the community ecology of plant microbiota.

## Acknowledgements

We thank Total Grower Services for taking and sending the avocado and banana samples.

## Supplementary Figures

Figure S1. Representation of the experimental design. Samples from avocado or banana plants were split into root and leaf compartments. Each sample was cultivated en masse on R2A media. The total community and cultivated versions of the community were transplanted to a surrogate host (tomato, or ryegrass). After four or eight weeks, four replicate surrogate plants were harvested for each treatment. Samples were transplanted root to root or leaf to leaf.

Figure S2. Schematic representation of the mesocosm system. Pots had two layers of plastic to minimise dispersal: a gamma-irradiated layer encased the pots (A) to prevent root and soil movement (C), and, plastic sleeves protruded into the air but remained opened at the top to prevent high humidity (B) to avoid aerosol and shoot-to-shoot contamination (C).

Figure S3. The % of taxa whose 95% credible region is above, or below, zero across root and leaf taxa for each factor and interaction term. Here plant refers to the surrogate or recipient plant species and donor species refers to the donor plant species

Table S1. Anova results from the four ratio metrics.

## Notes

https://github.com/ch16S/host_to_host_transplant

### Competing Interest Statement

Neil Wilson and Chris Baldock own shares in Metagen PTY LTD

