## Supplementary figures and images for "Evaluating host to host transplants as a method to study plant bacterial assembly"

### Figure S1

#### Slide 1
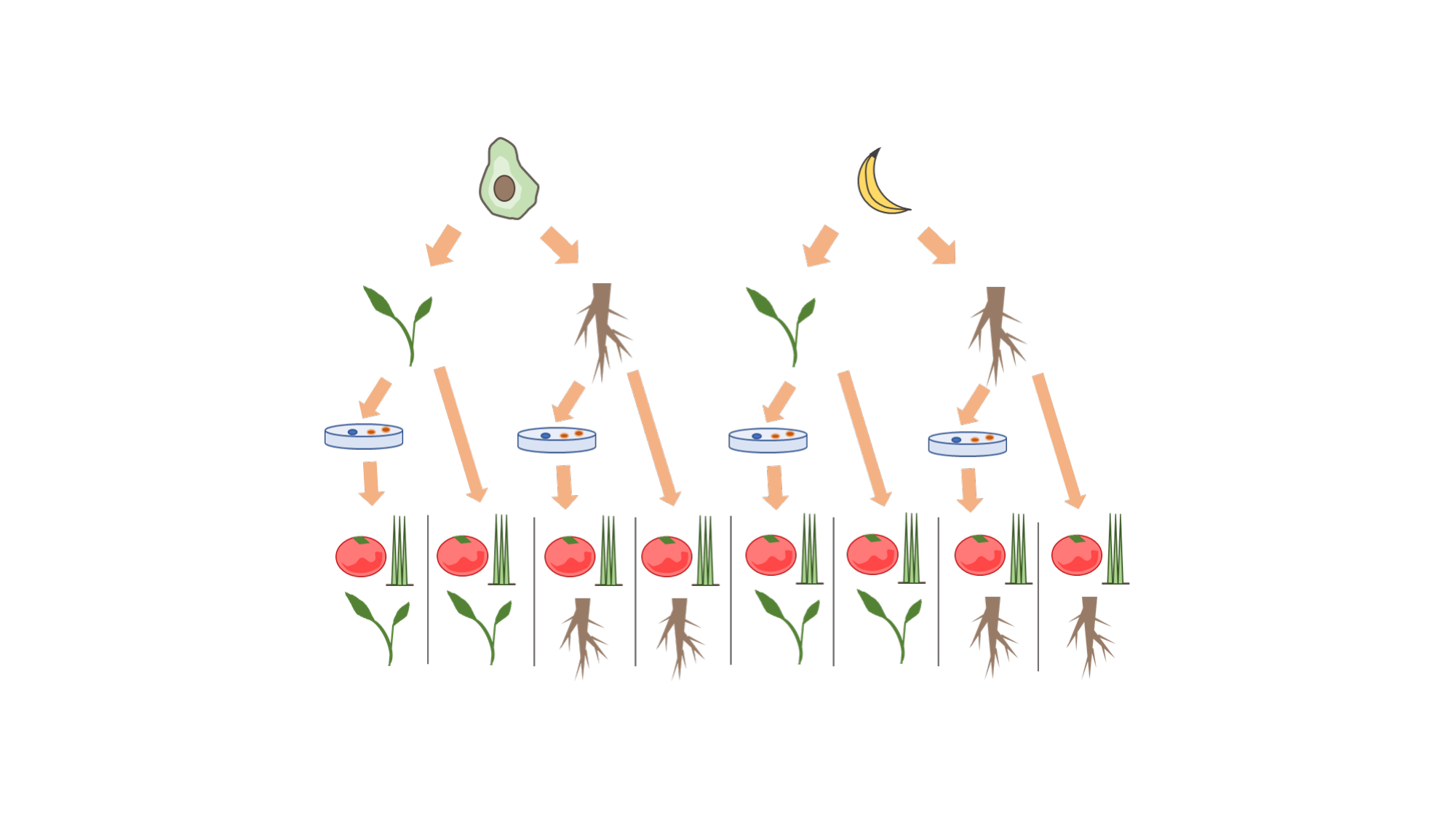

### Figure S2

#### Slide 1
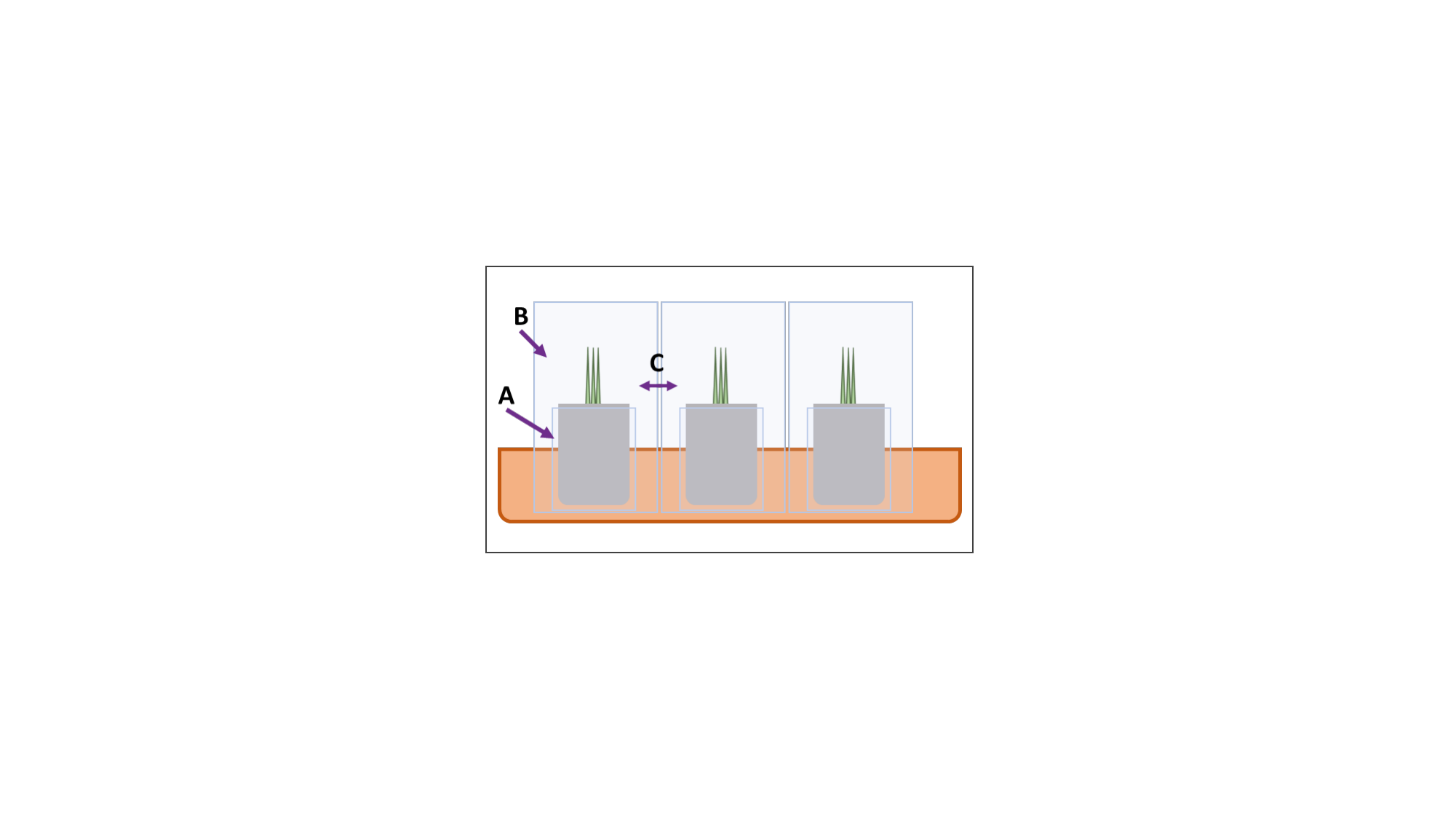

### Figure S3

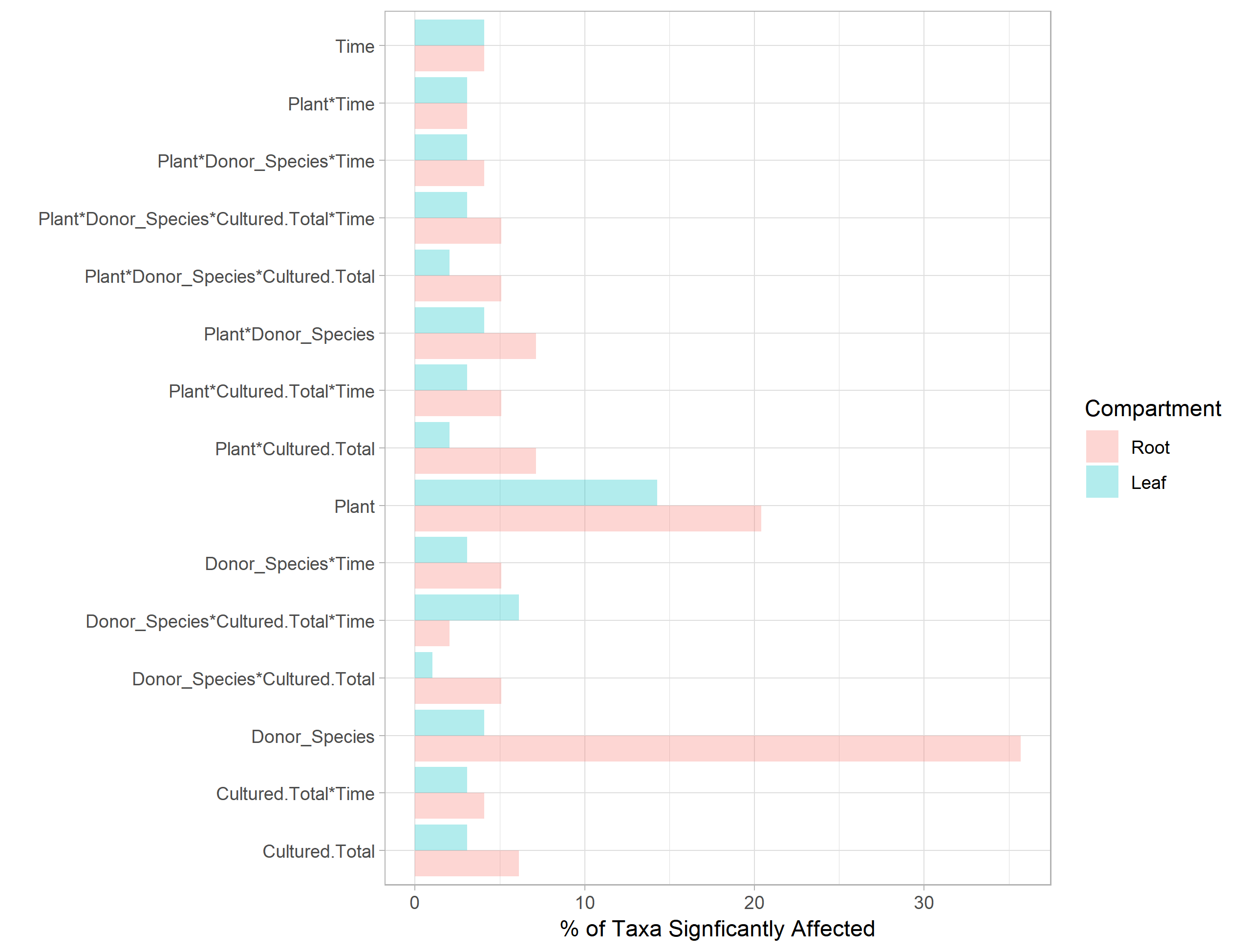
